# Improving dictionary-based named entity recognition with deep learning

**DOI:** 10.1101/2023.12.10.570777

**Authors:** Katerina Nastou, Mikaela Koutrouli, Sampo Pyysalo, Lars Juhl Jensen

## Abstract

**Motivation:** Dictionary-based named entity recognition (NER) allows terms to be detected in a corpus and normalized to biomedical databases and ontologies. However, adaptation to different entity types requires new high-quality dictionaries and associated lists of blocked names for each type. The latter are so far created by identifying cases that cause many false positives through manual inspection of individual names, a process that scales poorly.

**Results:** In this work we aim to improve block lists by automatically identifying names to block, based on the context in which they appear. By comparing results of three well-established biomedical NER methods, we generated a dataset of over 12.5 million text spans where the methods agree on the boundaries and type of entity tagged. These were used to generate positive and negative examples of contexts for four entity types (genes, diseases, species, chemicals), which were used to train a Transformer-based model (BioBERT) to perform entity type classification. Application of the best model (F1-score=96.7%) allowed us to generate a list of problematic names that should be blocked. Introducing this into our system doubled the size of the previous list of corpus-wide blocked names. Additionally, we generated a document-specific list that allows ambiguous names to be blocked in specific documents. These changes boosted text mining precision by *∼*5.5% on average, and over 8.5% for chemical and 7.5% for gene names, positively affecting several biological databases utilizing this NER system, like the STRING database, with only a minor drop in recall (0.6%).

**Availability:** All resources are available through Zenodo https://doi.org/10.5281/zenodo.10800530 and GitHub https://doi.org/10.5281/zenodo.10289360.

## Introduction

Named entity recognition (NER) is an important task in text mining that allows the identification of domain-specific terms or phrases – also known as named entities – in text and their classification into entity types [Nadeau and Sekine, 2007]. In the biomedical domain, named entities include genes, chemicals and diseases among others. NER is a prerequisite for the performance of other text mining tasks that allow the extraction of valuable information from text, such as relation extraction (RE) [Perera et al., 2020].

Dictionary-based methods, which leverage pre-defined dictionaries compiled from extensive collections of terms for each entity type, have been a prominent approach in NER [Leser and Hakenberg, 2005], as they simultaneously allow both the recognition and normalization of identified names in text. However, constructing dictionaries tailored for biomedical text mining demands significant time and expert knowledge, posing challenges in achieving optimal results [Wang et al., 2018].

We have in the past developed a robust, fast dictionary-based tagging engine, the JensenLab `tagger`, which serves as the core of an NER system that is applicable to many biomedical problems [Jensen, 2016]. The software can process thousands of PubMed abstracts per second and this high speed makes it well-suited for both real-time text mining [Pafilis et al., 2016] and NER in massive text corpora [Westergaard et al., 2018]. Combining this tagger with other dictionaries forms the basis for extraction of associations among genes/proteins [Szklarczyk et al., 2023], small molecule compounds [Szklarczyk et al., 2016], cellular components [Binder et al., 2014], tissues [Palasca et al., 2018], and diseases [Grissa et al., 2022], among others. Text-mining results obtained from these applications are normalized to identifiers from suitable databases and ontologies [Schriml et al., 2012, Wang et al., 2009, Maglott et al., 2000, Hubbard et al., 2002], and are used in several biological databases, which are freely available as a suite of web resources. These databases include the STRING database of protein interactions [Szklarczyk et al., 2023].

Despite its simplicity, dictionary-based NER is a powerful method that can perform surprisingly well in the biomedical domain. While software does matter, the most important part of any dictionary-based NER system is the dictionary itself. Adaptation of the NER system to a different domain requires the construction of a new high-quality dictionary and an associated block list of names. The manual creation of block lists for `tagger` was so far done through manual inspection of the names that occur most frequently in a large text corpus to identify those that would produce numerous false positives and should be disregarded. This process scales poorly, due to the manual labor involved.

The majority of the biomedical text mining community has been using deep learning-based and specifically Transformer-based methods [Miranda-Escalada et al., 2023] to perform several tasks, including NER, RE and Question-Answering. Moreover, Large Language Models have recently emerged as an alternative also for tasks other than Question-Answering [Wang et al., 2023]. We wanted to explore how we could take advantage of deep learning-based methods to improve the JensenLab suite of web resources. The intrinsic nature of these resources necessitates named entity normalization in conjunction with NER, and a fast and lightweight tagging system allowing weekly updates. It is prohibiting in terms of cost to perform such updates using deep-learning methods. Moreover, given the lack of suitable corpora to train deep learning-based methods for named entity recognition and normalization across all entity types, our objective did not center around devising an entirely new method. Rather, we could see a prospect in enhancing the precision of our dictionary-based NER system through the incorporation of state-of-the-art methods.

In this study, we present an alternative method for block list generation, which can bring many of the benefits of deep learning to dictionary-based NER, without sacrificing the advantages of the latter. To address the limitations of manual block list creation, we automated this crucial step by unifying and comparing results of three biomedical NER methods. We sought to create a large comprehensive dataset of text spans where different NER methods agreed on entity boundaries and types. This consensus dataset served as the foundation for generating both positive and negative examples for four entity types: genes or gene products, diseases, species, and chemicals. We then trained a Transformer-based model, specifically BioBERT [Lee et al., 2019], to perform entity type classification using that dataset. Ultimately we employed this model to accurately identify problematic names for blocking, resulting in a substantial expansion of the existing block list, ensuring a precision enhancement without compromising recall. Additionally, we created document-specific lists to address ambiguous name occurrences within specific documents. The successful implementation of these strategies significantly enhanced the quality of all web resources populated with associations generated by `tagger`. Moreover, this work serves as a guide for automating the identification of names to be blocked in various dictionary-based text mining systems, setting a precedent for more efficient and accurate NER in the biomedical domain.

## Materials and methods

### The dictionary-based tagger

As mentioned above text mining is integral to a suite of web resources with associations among biomedical entities [Jensen, 2016]. The initial step in extracting these associations from the literature involves the recognition of biomedical entities by the `tagger`, laying the foundation for relation extraction (RE). It is evident that the quality of the NER system directly impacts the quality of the generated associations.

Blocking and allowing names for the `tagger` is a key step to ensuring good quality. It operates across two levels: global (i.e. corpus-wide) and local (i.e. document-specific), utilizing two distinct methods for incorporating terms: automatic and manual additions. The generation of manual lists involves meticulous inspection of the most frequent names in the literature before addition, while that of automatic lists does not involve any human intervention. Both lists are combined to create a final list, favoring the manually curated list in case of conflicting overlaps between the two.

Operations on the global level involve both global blocking to identify names generally unsuitable for text mining, such as “Bad” or “ask”, which should be altogether removed for increased precision, and global allowing to safeguard names verified and accurately labeled as biomedical entity names of the correct type (e.g. “ATM”). Furthermore, the globally curated allowing functionality acts as a protection against potential inaccuracies from an auto-generated global block list. In local contexts, blocking or allowing decisions are tailored to the document at hand. For instance, “wingless” predominantly signifies a protein, yet in select instances, it refers to apterous insects, and local blocking in these instances is preferred to incorrect tagging. Similarly, a name like “toy”, which should be blocked globally, might be allowed in a specific document where it refers to the transcription factor “toy”. Despite the potential for both manual and automatic approaches in local contexts, considering the amount of effort these require, only automatic additions of names to the local list make sense.

The `tagger` functions in a case-insensitive manner and allows for arbitrary insertion or deletion of spaces and hyphens in the names and punctuation characters before or after them, both important for capturing the orthographic variation of biomedical named entities [Pafilis et al., 2013]. By contrast, all lists mentioned above function in a case-sensitive manner, affecting only specific versions of a name when added therein (e.g. adding “bad” will not affect the tagging of “BAD”).

During dictionary generation, clashes may arise, such as “NO” existing as a synonym in both the gene and chemical dictionaries. To resolve these clashes, a preference hierarchy based on source quality is implemented during the generation of the dictionaries for `tagger`. Diseases hold precedence over species, followed by genes or gene products, and finally chemicals. This hierarchy dictates that in the example above, “NO” will be tagged as a gene unless otherwise filtered. The resolution of such conflicts necessitates specific entity type filtering during dictionary creation. By employing filters, conflicts like overlapping gene and chemical names for “NO” can be addressed, ensuring accurate tagging.

The trust hierarchy in dictionaries and the inherent complexity of manually integrating terms for resolving dictionary clashes emphasize the significance of automatically blocking names after dictionaries have been built. This automatic process stands as the primary approach for enhancing the precision of `tagger` results, presenting a streamlined and effective means for optimizing accuracy for both NER and RE. For more details on dictionary generation for the `tagger`, please refer to [Jensen, 2016].

### Consensus dataset

The first step towards creating a method for automatically detecting problematic names in a biomedical dictionary is to identify high-quality examples to train the method on. These come in the form of a labeled dataset that allows supervised training for the task of biomedical entity type classification. To ensure that such a dataset has high coverage, a very large pool of both positive and negative examples is required. Since obtaining enough examples of properly labeled named entities via manual annotation is prohibitive — due to the time and resource demands of the effort — we opted for an automatic approach, which can provide a dataset that is simultaneously large and of high quality. To this end, we compared the NER results of PubTator [Wei et al., 2019], EVEX [Van Landeghem et al., 2013] and `tagger` [Jensen, 2016], thus identifying millions of text spans where all three methods agree on the boundaries and the type of entity tagged.

As mentioned, `tagger` is a dictionary-based method, with a dictionary based on Ensembl [Hubbard et al., 2002], Refseq [Maglott et al., 2000], PubChem [Wang et al., 2009], NCBI taxonomy [Schoch et al., 2020], and Disease Ontology [Schriml et al., 2012]. On the other hand, PubTator encompasses different text-mining tools [Huang et al., 2011, Wei et al., 2012, Leaman et al., 2013] to detect biomedical entities following different nomenclatures than `tagger`, while EVEX uses the Turku Event Extraction System [Björne et al., 2010] together with BANNER [Leaman and Gonzalez, 2008] and the McClosky-Charniak Parser [McClosky and Charniak, 2008]. The selection of the three systems is appropriate for creating a highly precise consensus dataset of training examples for genes or gene product (ggp), disease (dis), species (org), and chemical (che) named entities, as these methods rely on different architectures and/or nomenclatures to produce NER results for the four entity types they have in common.

To train a model for classifying entities, we designed a dataset consisting of biomedical entities within their textual context, all labeled with the biomedical entity’s type. To achieve this, we constructed positive examples using the consensus results of all three aforementioned NER methods, i.e. matches where all three methods agree on boundaries and entity type, comprising 200-word contexts around instances of the four designated entity types (100 words to the left and right of each entity mention). At the same time, we compiled a negative set from contexts around nouns and noun phrases where none of the methods produced a match corresponding to any of the targeted named entity types (neg class).

### Experimental setup

The task of entity type classification has clear analogies to NER, an area where the current state-of-the-art methods predominantly utilize models based on the Transformer architecture [Vaswani et al., 2017]. These models are initially pre-trained on large collections of text to produce a general language model, and can later be fine-tuned to perform specific tasks, such as NER. We have selected the cased BioBERT base model v1.1 [Lee et al., 2019], a biomedical domain-specific pre-trained deep language representation model based on BERT [Devlin et al., 2018], to train a multi-class entity type classifier on the consensus dataset described above.

BERT-based models are pre-trained using masked language modeling [Devlin et al., 2018], i.e. hiding certain words with a special mask token and asking the model to fill in the gaps. We have leveraged this feature of BioBERT to train our classifier to learn only how the contexts around each biomedical entity type look, without access to the text of the named entity mentions themselves. Specifically, we masked all entity mentions of interest in the consensus set, and set the label for each example based on the masked entity, e.g. “*Steroid excretion in patients receiving [MASK] compounds*.” would get a *chemical* label, while “*When [MASK] becomes mutated, it loses its function*.” would get a *gene or gene product* label. We have generated two datasets with the format described above: a 125k dataset with 125,000 training examples and a 12.5M dataset of 12,500,000 training examples.

During fine-tuning, we updated the weights of the pre-trained BioBERT model to classify the examples described above as belonging to one of the five training classes (ggp, org, dis, che, neg) with the output softmax layer configured to transform the contextualized representations generated by the BERT model into segment-level probabilities for each class. These probabilities were derived by aggregating information from tokens within each segment, enabling the prediction of the most likely class for the entire segment. To select the best model, we tuned the hyperparameters of the BioBERT model through a grid search to find the best combination of values for the learning rate (5e-5, 3e-5, 2e-5, 1e-5, and 5e-6), mini batch size (32 and 64), number of epochs (2, 3, and 4), maximum sequence length (256), and context size (200). The values in parentheses for each parameter were those suggested in the BioBERT publication and some extensions to those, based on our initial experiments. For more details on the experimental setup please refer to our code base, available through GitHub. The experiments were repeated four times and the results are expressed as a mean and standard deviation of the F1-score (micro-averaged over all labels in the sets). The abovementioned experiments were run using the 125k dataset described above by assigning 125,000 examples to the training set and 62,500 to the development set. The hyperparameters producing the best total mean F1-score on the development set were selected for training a model on the 12.5M dataset — 12.5 million examples were used for training and 62,500 to calculate the test set performance on this set.

### Block List generation and prediction

To generate automated block lists, we used all PubMed abstracts (as of August 2022) and all full-texts available in the PMC BioC text mining collection [Comeau et al., 2019] (as of April 2022). We then ran `tagger` using the manually curated block lists only. We generated examples for all contexts surrounding the `tagger` matches of the four positive classes, and we used the best-performing model for entity type prediction on these examples.

As explained above, for each example, the classifier produces a probabilistic score of belonging to each class (including the negative class) and, based on these, we can calculate the score of not belonging to the `tagger`-assigned class P(*¬*C), where C stands for class. To generate a corpus-wide block list, we do an unweighted score average of all P(*¬*C) within each document for each name. We then average these document-level scores across the entire corpus — considering that matches in different documents are independent — and generate a single score per name of not belonging to the `tagger`-assigned class across the entire literature. To generate the global block list we had an additional requirement, that there are at least two documents with one mention for each name. Afterwards, we manually inspected the lists to identify the thresholds that we would use to add names to the global block list, since those could be different for each class.

The fact that we had a probabilistic score for each match allowed us to explore the use of local (document-level) predictions to resolve ambiguous names, by aggregating within-document scores for each name. This allowed us to test both the generation of lists of terms that could be locally blocked or allowed — the latter only for terms that are automatically blocked globally. Once again, the scores that would serve as thresholds for inclusion in these lists had to be manually identified by inspecting random samples of the lists. To identify names that would be locally allowed we calculated a ratio based on the formula below:

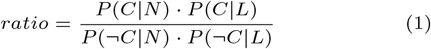

where C stands for class and corresponds to the five entity types used to train the classifier (gene or gene product, disease, species, chemical and negative), P(C|N) is the global probabilistic score of a term belonging to that specific class, P(C|L) is the local probabilistic score estimate for the term belonging to that class, and *¬*C is “not C”, so P(*¬*C|N) = (1 *−* P(C|N)) and P(*¬*C|L) = (1 *−* P(C|L)). This formula allows us to override even very high global confidence scores when trying to add names to the local allow list. The calculation of the ratio for identifying names to be locally blocked is analogous to (1).

### Evaluation

To evaluate the effects of block lists we generated four sets of results by performing four different `tagger` runs:

1. run with a combined automatically generated and manually curated block list (***curated+auto***): This run involved using a combination of both automatically generated and manually curated block lists for `tagger` and resembles what is currently done for JensenLab web resources.
2. run with manually curated block lists (***curated only***): in this run, only the manually curated block list was used.
3. run with automatically generated block lists (***auto only***): in this run we only used block lists that were automatically generated from the fine-tuned BioBERT model. To generate these lists, we had to run `tagger` without any block list (only the built-in `tagger` regular expressions were used to block names) and we repeated the process for block list generation, using the thresholds that we determined from the process described in the previous section.
4. run without any block list (***no block list***): this involved running `tagger` without utilizing any block list. We also adapted the tagger code to exclude the use of built-in regular expressions. The adapted code is provided through Zenodo.

We conducted evaluations using two distinct approaches. Firstly, since the main incentive for updating the block lists is improving the JensenLab suite of web resources, we aimed to assess the impact of block lists in co-occurrence-based relation extraction. Specifically, we evaluated the results from the four runs above against two datasets: the KEGG database [Kanehisa and Goto, 2000] gold standard used for evaluations by the STRING database [Szklarczyk et al., 2023], and the gold standard of protein-disease associations used by the DISEASES database [Grissa et al., 2022].

Secondly, to evaluate the direct impact of automatically adding names in the block lists on NER, we manually checked 500 random matches of each class from the ***curated only*** run and calculated `tagger`’s precision for each class using the formula below:

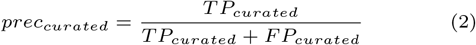

where TP_curated_ is the number of true positive matches, i.e. those belonging to the class assigned by the `tagger`, and FP_curated_ is the number of false positive matches.

To assess the effect of the addition of the automatic block list, we checked 200 matches per class that got removed as a result of incorporating the automatically generated block list. For this purpose, we compared the results of the ***curated only*** run and the ***curated+auto*** run and identified the set of matches that got removed between these two runs. Then we randomly selected a subset of these removed matches to inspect, and calculated the precision of automatic blocking using the formula below:

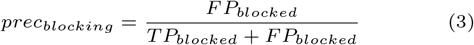

The prec_blocking_ is the fraction of automatically blocked names that were correctly blocked, i.e. former FPs that were removed by blocking.

To calculate the effect of automatic blocking on precision and recall, we need to introduce weighted TPs and FPs:

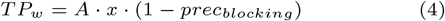

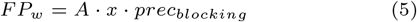

where *A* = 500 (*TP*_*curated*_ + *FP*_*curated*_) and *x* is the class-specific automatic block rate (calculated from the difference in the number of matches between the ***curated only*** and ***curated+auto*** runs). From these, the precision using both curated and automatic block list (*prec*_*c*+*a*_) and the relative difference in recall (*rec*_*diff*_) for a given class can be calculated as:

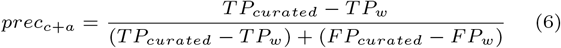

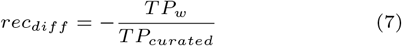

## Results and discussion

### Dataset generation and model training

By obtaining the consensus results of `tagger`, PubTator, and EVEX, we generated a dataset with millions of text spans of positive and negative examples, which we split into a 125k and a 12.5M set as described in 2.3. The number of examples was balanced between the 5 classes, as shown in Table 1 for the training datasets. A manual inspection of 500 names (100 of each class) from this dataset allowed us to calculate its precision, which was found to be 99.8% (Supplementary Table 1, available through Zenodo), showcasing the high quality of this dataset.

**Table 1.** Number of examples for each class in the 12.5M and 125k training sets

| Entity type | Number of examples |  |
| --- | --- | --- |
|  | 12.5M set | 125k set |
| Gene or gene product | 2,292,475 | 23,105 |
| Chemical | 3,705,213 | 37,205 |
| Disease | 2,083,848 | 20,795 |
| Species | 1,918,464 | 19,402 |
| Negative | 2,500,000 | 24,493 |
| Total | 12,500,000 | 125,000 |

Our hyperparameter grid search on the 125k set revealed that the BioBERT_base_ model with a learning rate of 2^*−*5^ and a batch size of 32 was the best model trained on the set of 125,000 examples, with a mean F-score of 94.83% (SD=0.0356%) on the development set. All models in this grid had a mean F-score of 94% or above. Detailed results are presented in Supplementary Table 2. We used the set of hyperparameters above to train a model on the 12.5M set for one epoch. This model achieved an accuracy of 96.67% (SD=0%) on the test set of 62,500 examples and is the one that is further used for all prediction runs. The fine-tuned Tensorflow model on the 12.5M set is available through Zenodo.

### Block List generation

The `tagger` was executed using the full `tagger` dictionary and only manually curated lists on a corpus of the entire biomedical scientific literature as of August 2022, which consisted of 34,420,049 documents. The dictionary and block list files, as well as the input documents used in this run are available via Zenodo. There were a total of 75,419,774 gene or gene product, 78,615,064 chemical, 39,995,175 disease, and 62,921,879 species matches as a result of that run. For each match, a 200-word context was generated from the input document and classified using the fine-tuned model for entity type prediction.

Using the process detailed in 2.4, we generated an initial set of global block lists to identify the probabilistic score threshold to use for each entity type. These were chosen empirically based on the relative densities of true and false positives at threshold values for P(*¬*C) in steps of 0.05 starting from 0.5. A spreadsheet with the lists of names checked during this process is available through Zenodo. The best threshold choice for genes or gene products, species, and chemicals was 0.5, while for diseases the value was 0.85.

We also had to manually determine the ratios for local allow/block lists, by manually inspecting a random set of blocked matches within their textual context. This inspection showed that the generation of good quality lists for diseases and species was not possible. So we generated lists only for genes or gene products and chemicals, using a ratio (Equation 1) of 1000 as the threshold of inclusion. As for the local allow lists genes or gene products was the only class with good quality lists and the ratio for those was set to a very high value (10^15^).

Introducing the lists above into the `tagger` dictionary approximately doubled the size of the global list from 148,143 names to 350,862 names. The local (document-specific) allow/block list was introduced for the first time in the text-mining system and contains 155,660 names blocked and 9,611 names allowed in 144,585 and 9,458 documents, respectively. This led to a block rate of 7.4%, leading to a total reduction of 51,675,498 unique `tagger` matches in the four targeted classes due to the additional names that were blocked (from 693,282,180 matches to 641,606,682 matches). Specifically, for chemicals the block rate is *∼*11% (24,026,228 matches removed), *∼*9% for genes or gene products (20,551,927 matches removed), *∼*4% for species (4,897,488 matches removed), and *∼*2% for diseases (2,199,855 matches removed).

### Evaluation

To measure the effect of introducing automatically generated block lists on `tagger`’s performance, we run `tagger` using four different setups as described in Methods. We detected all co-occurring protein–protein and protein–disease pairs in the literature resulting from these four runs and evaluated the performance of co-occurrence-based relation extraction using a functional protein–protein association gold standard from KEGG [Kanehisa and Goto, 2000] and a protein–disease gold standard from DISEASES [Grissa et al., 2022]. The results from this evaluation are shown in Figure 1.

**Fig. 1.**
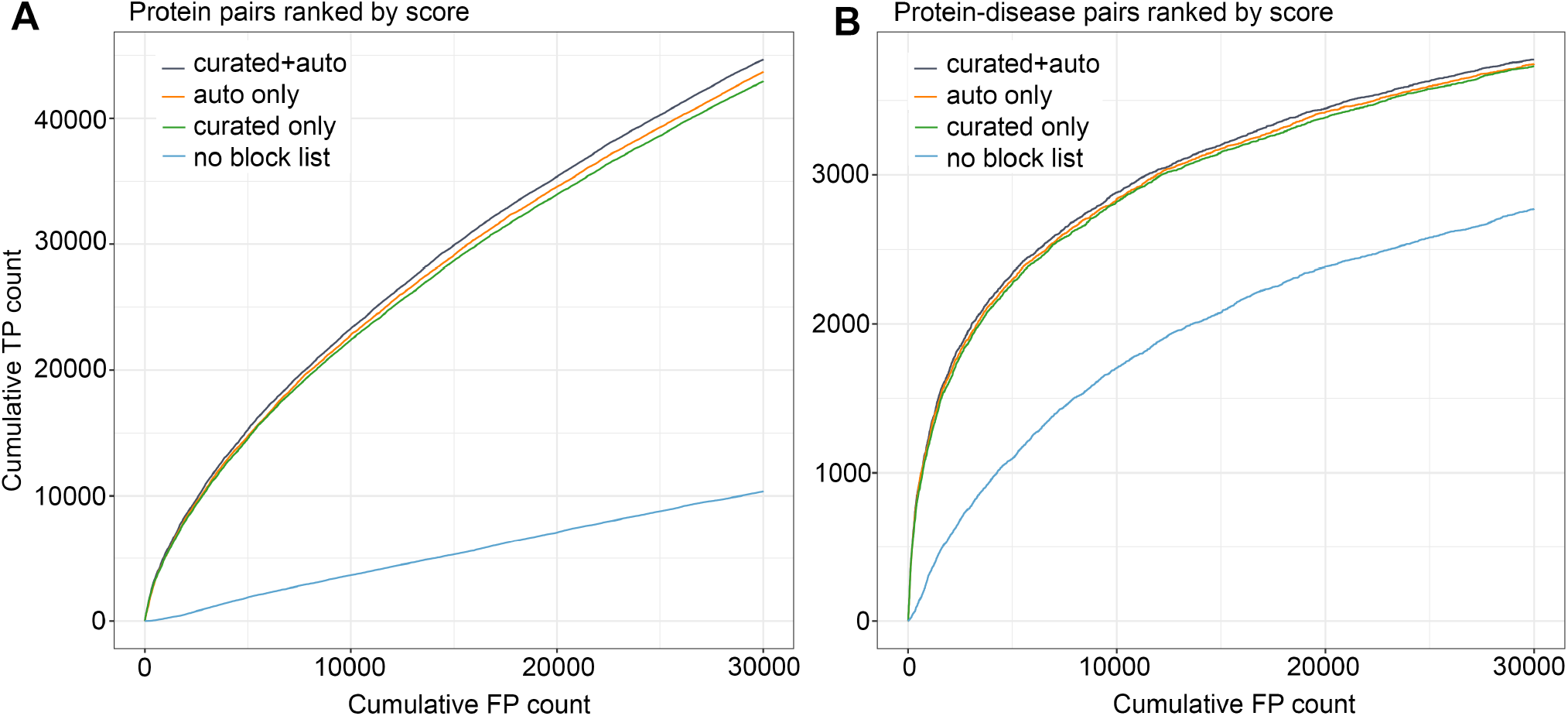
Cumulative True Positive (TP) /False Positive (FP) curves to evaluate human (A) functional protein-protein associations and (B) protein– disease associations. The pairs are ranked based on their significance score. For details on the calculations of this score please refer to [Franceschini et al., 2012]. The curves are showing only up to Cumulative FP count of 30,000 pairs.

Figure 1 shows that the incorporation of any block list in `tagger` markedly enhances the quality of its generated results. Notably, best results are achieved with our newly proposed method, i.e. by the combination of the automatically generated and the curated block lists. However, the differences between the three block list-including runs are not particularly pronounced. Even when focusing on the initial segments of the curves where more cumulative true positives are expected, the differences are still slim. A potential explanation for this result is that proteins present in the two gold standard datasets tend to be well-studied. This means that they will frequently show up in the literature and that problematic names are thus relatively easy to find both by manual curation (green line) and by the automatic approach (orange line). Consequently, these datasets may not accurately reflect the potential benefits derived from the automated blocking of names, as manual curation will almost certainly give worse coverage of the less studied proteins, which are not in these gold standards. We thus opted to employ a second strategy.

For the second strategy, we assessed the quality of automatically blocked names by manually inspecting them. Table 2 shows the precision of `tagger` per class when using only the curated block list and the effect of automatic blocking of names, allowing us to see the changes in the dictionary-based system’s precision and recall. To calculate the initial precision of `tagger` for each class we have selected 500 matches from the **curated only** run for each of the four targeted entity types, did a manual evaluation of the correct assignment to this class by tagger and assigned a TP or FP label (Supplementary Tables 3– 6, available through Zenodo). To assess if a match is a TP or FP, we manually check the names within the specified context (i.e. within the documents in which the matches appear), to assess if they belong to their assigned class. Correct normalization was not a requirement to assign a TP label. The precision of `tagger` when run using only the curated list is calculated using formula 2 and is presented in the first column of Table 3.

**Table 2.** Precision and Relative Recall calculations for `tagger`

| Entity type | precision<br>curated | precision<br>curated<br>+auto | precision<br>difference | recall<br>difference |
| --- | --- | --- | --- | --- |
| Gene or<br>gene product | 89.8% | 97.6% | +7.8% | -1.1% |
| Chemical | 76.2% | 85.1% | +8.9% | -0.6% |
| Disease | 98.4% | 99.8% | +1.4% | -0.6% |
| Species | 88.4% | 91.8% | +3.4% | -0.3% |
| Average | 88.2% | 93.6% | +5.4% | -0.6% |

Then we randomly selected 200 matches for each class, that were removed due to the introduction of the automatically generated block list and manually checked to evaluate which ones were TPs and which were FPs (Supplementary Tables 7–10). We used formula 6 to calculate the precision after the introduction of this block list. Moreover, we have used formula 7 to calculate the relative difference in recall due to the addition of the automatically generated block list.

Overall, one can see that while the introduction of the automatically generated block lists will inadvertently result in a recall reduction — due to the blocking of some good names — the precision gain clearly overshadows the former. The effect is more evident for classes where dictionary-based NER faces more problems, like chemicals, and very slight in cases where dictionary-based text mining already performs very well, like diseases. These results support our decision to introduce the automatically generated block lists in `tagger`’s dictionaries and have a positive effect for JensenLab web resources, including the latest version of the STRING database. Additionally, for evaluating the effect of the new block list on dictionary-based species NER, we used the recently published S1000 corpus [Luoma et al., 2023]. The F-score increased by 1% (from 84.7% to 85.7%) in this dataset, which was a result of a 2.6% increase in precision and a 0.4% decrease in recall, showcasing an overall positive effect.

## Conclusions

In this work, we propose a workflow designed to automate the process of adding names in a block list for dictionary-based NER systems and use the `tagger` to assess the results of this process. We have generated and used a very large dataset of 12.5 million text spans, to generate positive and negative examples of contexts for four entity types, namely genes or gene products, diseases, species and chemicals, and a negative class. By training a highly accurate (F-score=96.7%) Transformer-based entity type classifier on this dataset, and using it to do entity type prediction on all `tagger` matches resulting from a run with a curated block list, we have managed to effectively double the size of the global block list that `tagger` uses, which would not have been feasible by manual curation. At the same time, we have incorporated for the first time an extensive automated local allow/block list which supports the document-specific resolution of ambiguous names.

Evaluation of the newly generated lists demonstrated positive effects on both dictionary-based NER and RE. Subsequently, these enhanced lists have been directly incorporated into `tagger`, without altering how the tool works or adversely affecting its speed. The improved dictionaries and annotation results are thus made publicly available immediately, through the RESTful APIs that serve the web resources with text mining results produced by the `tagger`, including the widely used STRING database. This integration ensures immediate access to improved resources, advancing the utility and accuracy of `tagger` in several biomedical text mining applications.

## Acknowledgements

We thank the CSC – IT Center for Science for generous computational resources. We would like to thank Rebecca Kirsch for providing the code to generate Figure 1.

## References

J. X. Binder, S. Pletscher-Frankild, K. Tsafou, C. Stolte, S. I. O’Donoghue, R. Schneider, and L. J. Jensen. Compartments: unification and visualization of protein subcellular localization evidence. Database, 2014, 2014.

J. Björne, F. Ginter, S. Pyysalo, J. Tsujii, and T. Salakoski. Complex event extraction at pubmed scale. Bioinformatics, 26(12):i382–i390, 2010.

Comeau, C. Wei, R. Islamaj Doğan, and Z. Lu. Pmc text mining subset in bioc: about three million full-text articles and growing. Bioinformatics, 35(18):3533–3535, 2019.

J. Devlin, M.-W. Chang, K. Lee, and K. Toutanova. Bert: Pre-training of deep bidirectional transformers for language understanding. arXiv, 2018.

Franceschini, D. Szklarczyk, S. Frankild, M. Kuhn, M. Simonovic, A. Roth, J. Lin, P. Minguez, P. Bork, Von Mering, et al. String v9. 1: protein-protein interaction networks, with increased coverage and integration. Nucleic acids research, 41(D1):D808–D815, 2012.

Grissa, A. Junge, T. I. Oprea, and L. J. Jensen. Diseases 2.0: a weekly updated database of disease–gene associations from text mining and data integration. Database, 2022:baac019, 2022.

M. Huang, J. Liu, and X. Zhu. Genetukit: a software for document-level gene normalization. Bioinformatics, 27(7): 1032–1033, 2011.

T. Hubbard, D. Barker, E. Birney, G. Cameron, Y. Chen, L. Clark, T. Cox, J. Cuff, V. Curwen, T. Down, et al. The ensembl genome database project. Nucleic Acids Res., 30 (1):38–41, 2002.

L. J. Jensen. One tagger, many uses: Illustrating the power of ontologies in dictionary-based named entity recognition. bioRxiv, page 067132, 2016.

M. Kanehisa and S. Goto. Kegg: kyoto encyclopedia of genes and genomes. Nucleic Acids Res., 28(1):27–30, 2000.

R. Leaman and G. Gonzalez. Banner: an executable survey of advances in biomedical named entity recognition. In Biocomputing, pages 652–663. World Scientific, 2008.

R. Leaman, R. Islamaj Doğan, and Z. Lu. Dnorm: disease name normalization with pairwise learning to rank. Bioinformatics, 29(22):2909–2917, 2013.

J. Lee, W. Yoon, S. Kim, D. Kim, S. Kim, C. H. So, and J. Kang. BioBERT: a pre-trained biomedical language representation model for biomedical text mining. Bioinformatics, 36(4):1234–1240, 2019.

U. Leser and J. Hakenberg. What makes a gene name? named entity recognition in the biomedical literature. Brief. Bioinform., 6(4):357–369, 2005.

J. Luoma, K. Nastou, T. Ohta, H. Toivonen, E. Pafilis, L. J. Jensen, and S. Pyysalo. S1000: A better taxonomic name corpus for biomedical information extraction. Bioinformatics, 39(6):btad369, 2023.

D. Maglott, K. Katz, H. Sicotte, and K. Pruitt. Ncbi’s locuslink and refseq. Nucleic acids research, 28(1):126–128, 2000.

D. McClosky and E. Charniak. Self-training for biomedical parsing. In Proceedings of ACL-08, pages 101–104, 2008.

A. Miranda-Escalada, F. Mehryary, J. Luoma, D. Estrada-Zavala, L. Gasco, S. Pyysalo, A. Valencia, and M. Krallinger. Overview of DrugProt task at BioCreative VII: data and methods for large-scale text mining and knowledge graph generation of heterogenous chemical–protein relations. Database, 2023:baad080, 2023.

D. Nadeau and S. Sekine. A survey of named entity recognition and classification. Ling. Invest., 30(1):3–26, 2007.

Pafilis, S. P. Frankild, L. Fanini, S. Faulwetter, C. Pavloudi, Vasileiadou, C. Arvanitidis, and L. J. Jensen. The species and organisms resources for fast and accurate identification of taxonomic names in text. PloS one, 8(6):e65390, 2013.

E. Pafilis, P. L. Buttigieg, B. Ferrell, E. Pereira, J. Schnetzer Arvanitidis, and L. J. Jensen. Extract: interactive extraction of environment metadata and term suggestion for metagenomic sample annotation. Database, 2016, 2016.

O. Palasca, A. Santos, C. Stolte, J. Gorodkin, and L. J. Jensen. Tissues 2.0: an integrative web resource on mammalian tissue expression. Database, 2018:bay003, 2018.

N. Perera, M. Dehmer, and F. Emmert-Streib. Named entity recognition and relation detection for biomedical information extraction. Front. cell dev. biol., page 673, 2020.

L. Schoch, S. Ciufo, M. Domrachev, C. L. Hotton, S. Kannan, R. Khovanskaya, D. Leipe, R. Mcveigh, O’Neill, B. Robbertse, et al. Ncbi taxonomy: a comprehensive update on curation, resources and tools. Database, 2020:baaa062, 2020.

M. Schriml, C. Arze, S. Nadendla, Y.-W. W. Chang Mazaitis, V. Felix, G. Feng, and W. A. Kibbe. Disease ontology: a backbone for disease semantic integration. Nucleic Acids Res., 40(D1):D940–D946, 2012.

D. Szklarczyk, A. Santos, C. Von Mering, L. J. Jensen, P. Bork, and M. Kuhn. Stitch 5: augmenting protein– chemical interaction networks with tissue and affinity data. Nucleic acids research, 44(D1):D380–D384, 2016.

D. Szklarczyk, R. Kirsch, M. Koutrouli, K. Nastou, Mehryary, R. Hachilif, A. L. Gable, T. Fang, N. T. Doncheva, S. Pyysalo, et al. The string database in 2023: protein–protein association networks and functional enrichment analyses for any sequenced genome of interest. Nucleic Acids Research, 51(D1):D638–D646, 2023.

S. Van Landeghem, J. Björne, C.-H. Wei, K. Hakala, S. Pyysalo, S. Ananiadou, H.-Y. Kao, Z. Lu, T. Salakoski, Y. Van de Peer, et al. Large-scale event extraction from literature with multi-level gene normalization. PloS one, 8(4):e55814, 2013.

Vaswani, N. Shazeer, N. Parmar, J. Uszkoreit, L. Jones, N. Gomez, L. Kaiser, and I. Polosukhin. Attention is all you need. In NIPS’17, NIPS’17, page 6000–6010, Red Hook, NY, USA, 2017. Curran Associates Inc.

S. Wang, X. Sun, X. Li, R. Ouyang, F. Wu, T. Zhang, J. Li, and G. Wang. Gpt-ner: Named entity recognition via large language models. arXiv, 2023.

X. Wang, C. Yang, and R. Guan. A comparative study for biomedical named entity recognition. IJMLC, 9(3):373–382, 2018.

Y. Wang, J. Xiao, T. O. Suzek, J. Zhang, J. Wang, and S. H. Bryant. Pubchem: a public information system for analyzing bioactivities of small molecules. Nucleic acids research, 37 (suppl 2):W623–W633, 2009.

Wei, H. Kao, and Z. Lu. Sr4gn: a species recognition software tool for gene normalization. PloS one, 7(6):e38460, 2012.

Wei, A. Allot, R. Leaman, and Z. Lu. Pubtator central: automated concept annotation for biomedical full text articles. Nucleic Acids Res., 47(W1):587–593, 2019.

Westergaard, H.-H. Stærfeldt, C. Tønsberg, L. J. Jensen, and S. Brunak. A comprehensive and quantitative comparison of text-mining in 15 million full-text articles versus their corresponding abstracts. PLoS comput. biol., 14(2):e1005962, 2018.

